# Exploration of the Timeliness of ICD Codes in Administrative Databases: A Nationwide Study

**DOI:** 10.1101/491951

**Authors:** Marie Simon, Bastien Rance, Sandrine Katsahian, Karim Bounebache, Grégoire Rey, Gilles Chatellier, Antoine Neuraz, Anita Burgun, Vincent Looten

## Abstract

**INTRODUCTION:** The ICD codes are ubiquitously available in hospital information systems and have been used in a number of areas such as epidemiology, phenotype-genotype association mining, surveillance of the use of drugs and medical devices and health care evaluation. We aimed to analyze the timeliness of the 3-character ICD-10 codes collected in the French national hospital discharge summary database between 2008 and 2017 and we classified the codes according to their evolution.

**MATERIAL AND METHODS:** We extracted all 3-character ICD-10 codes from all French hospital discharge summaries between 2008 and 2017. For each code and by the month, we computed a relative frequency; we also computed the overall amplitude of the study period. Temporal clustering, according to the SAX representation, was performed to classify the main evolution patterns.

**RESULTS:** We extracted 238,334,751 encounters corresponding to 56,621,773 distinct patients. 1,006 ICD codes presented a variation of the relative amplitude of frequencies lower than 50%, 510 codes between 50% and 100% and 521 greater than 100%. Out of the 2,037 codes included in the study, we kept the 1,758 for the temporal clustering. Four clusters were identified, including a global increase and a global decrease patterns.

**DISCUSSION:** The overall results showed a strong instability (i.e. large variation of frequency over time) of the use of ICD codes over time, with an important variation of the relative amplitude of the frequencies. We distinguished between external factors due to changes in billing, organization, policy or regulation and intrinsic factors due to epidemiological phenomenon. The detailed analysis of profiles show that the same cluster can contain profiles influenced by intrinsic epidemiological or external factors or both. Additional knowledge and sources are probably required to determine automatically the origin of the profile.

## 1 INTRODUCTION

The use of the diagnostic code for “*Malnutrition*” in the French national health administrative databases saw a six-fold increase between 2008 and 2017. Such finding is highly questionable, and researchers could reasonably wonder whether this increase denotes a real phenomenon in the French population or changes in data collection? Health administrative databases are massive repositories of data composed of medical procedures, prescriptions and diagnoses information[1,2] initially collected for administrative purposes. Therefore, the specifics of the data collection may vary depending on other incentives than epidemiology.

Secondary use of health administrative databases for research and public health has become more and more extensive because these databases exhibit many strengths: (i) they are population-based; (ii) data are collected as part of the routine care process, which makes investigations faster and research costs lower; (iii) the follow-up of specific populations can be traced for years or decades; (iv) the presence of several years of data allows for studying changes over time for numerous variables. Among these variables, diagnostic codes expresses as codes from the International Classification of Disease (ICD)provide standardized diagnoses information at local, national and international levels. In European countries, most hospitals leverage codes from the International Classification of Disease (ICD), version 10[3] for billing purposes. The ICD-10 coding system contains over 70,000 disease codes grouped into 22 higher order categories. These codes are ubiquitously available in hospital information systems and can be reused for research purposes[4]. ICD codes have been used in a number of areas such as epidemiology[5,6], phenotype-genotype association mining[7,8], distribution of risk factors and impact on major clinical outcomes[9–11], surveillance of the use of drugs and medical devices[12,13], health care evaluation and health economic evaluations[14]. In the context of large-scale collaboration at continental or international levels, like the Observational Health Data Sciences and Informatics (OHDSI) Network[15,16]), ICD data have been largely used to support federated analyses across institutions, e.g., to identify cancer patients[17], and across countries[18]). Over the last years there has been widespread development of medical data repositories, and the research community has put specific emphasis on enhancing the capacity of algorithms to automatically find and use the data, to analyze the data sets, and to mine the data for knowledge discovery[19]. However, there are several bottlenecks to consider when reusing health administrative data and a misinterpretation of the data can have dramatic consequences for research (and even public health decisions) and lead to invalid conclusions.

Regarding hospital data, Agniel *et al.*[20] note that EHRs capture more information than the simple results recorded. For example, the presence of a laboratory test order and the timing of when it was ordered, regardless of any other information about the test result, is indicative of “organizational" aspects such as, for example, the expertise of the clinicians. The presence of some tests may even have a significant association with the odds ratio of death. We hypothesized that, similarly to the data in the EHR, the disease codes in health administrative databases capture more information than the sheer frequency of the diseases in the population: the information carried by ICD codes in health administrative databases is impacted by intrinsic epidemiological factors, such as outbreaks or environmental events, and by external factors, such as financial or health policies. More precisely, we analyzed the whole set of ICD claim data collected in France between 2008 and 2017 to assess whether the temporal context of the variation of ICD codes could provide insight on the respective influence of epidemiological and external factors.

### 1.1 Related Works

Overtime the importance of epidemiological and external factors might evolve: a query built at a given time for patient selection may not be relevant at a later date, and lead to errors of interpretation. Mues *et al.*[21] discussed the influence of the Centers for Medicare and Medicaid Services policy on data collection and coding between 2005 and 2012. The authors show that the percentage of inpatient claims with a diagnosis code for coronary atherosclerosis suddenly increased in 2010 because of the expansion of the number of ICD-9/10 diagnosis and procedure code fields on a claim. Moreover, the number of ICD diagnosis codes expanded from 17,000 to 77,000 with the replacement of the ICD-9 by the ICD-10 in 2015. Sáez *et al.*[22] defined timeliness as “the degree of temporal stability of the data.” Rey *et al*.[23] proposed a statistical method that detects abrupt changes of the time course of mortality by cause measured with ICD codes. However, to the best of our knowledge, no study has proposed a model-free temporal clustering of ICD codes and a description of the evolution of ICD codes over time.

### 1.2 Scope and Objective

In this study, we aimed to analyze the timeliness of the 3-character ICD-10 codes collected in the French national hospital discharge summary database between 2008 and 2017. We classified the codes according to their evolution. We discussed the different classes to identify the importance of epidemiological and external effects.

## 2 MATERIAL AND METHODS

### 2.1 Study Design

We performed a retrospective study leveraging the French discharge summary database. We searched for the evolution over time of ICD-10 codes. We used the finest degree of granularity available for dates: namely the month.

### 2.2 Data source

The French National Health Data System (SNDS, for *Système National des Données de Santé*) is a national-level inclusive data repository that was implemented in 2017 to facilitate the secondary use of the French administrative databases. The SNDS includes the national hospitalization discharge summary database, covering 65 million people with a longitudinal and exhaustive follow-up of patients regarding their hospital’s history. The hospital discharge diagnoses have been recorded using ICD-10 since 2008[24]. We extracted all ICD-10 codes from all hospital discharge summaries between 2008 and 2017.

### 2.3 Definitions

We considered the 3-character ICD codes to reduce the number of codes while preserving a clinical or epidemiological interpretability. For example, the code C50.1 was standardized as C50. We extracted from the database tuples consisted of a timestamp (year-month), an ICD-10 code, the number of distinct patients annotated with the code for the given timestamp, the overall number of patients annotated with any code for the given timestamp. For each 3-character code and by the month, we computed the relative frequency, defined as follows: 
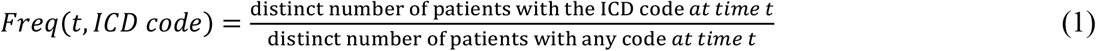

We applied a moving means, using the 6 months prior and after each point, to reduce the impact of extreme variations. To identify amplitude variations of the smoothed frequencies, we computed the relative amplitude defined as: 
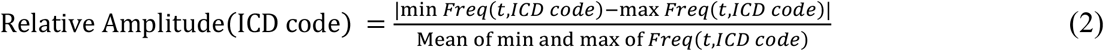

### 2.4 Temporal Clustering

We included in the temporal clustering analysis only ICD codes with data covering the entire period of the study (2008-2017) to exclude the infrequent codes. We performed a temporal hierarchical clustering analysis on the moving means time series. After normalizing the time series, we computed distances using the symbolic aggregate approximation (SAX) representation. Proposed by Lin *et al.*[26,27], the SAX representation is model-free and has been developed to deal with the heterogeneity of time series curves and pattern discovering. The SAX representation has two parameters: the amount of equal sized frames that the series will be reduced to (parameter w) and the size of the alphabet (parameter a, a>2). The SAX representation is performed in 2 steps: (1) the first step reduces the dimension of time series (2) the second step transforms the reduced time series into a symbolic representation. For the first step, each time series *C*_*raw*_ = (*C*_*raw*,1_,…,*C*_*raw,N*_) of length N (N=120 months) is represented in a w-dimensional space by a vector *C*_*SAX*_ = (*C*_*SAX*,1_,…*C*_*SAX,w*_) defined as: for *i* ∈ ⟦1,*w*⟧, 
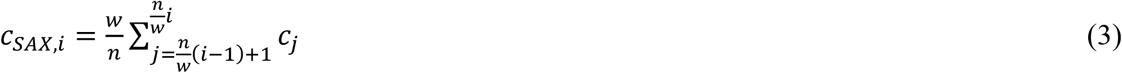

The second step transforms the vector *C*_*SAX*_ into a vector 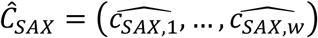 where 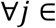, 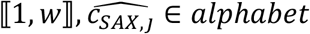, *alphabet* is the chosen alphabet (*Card*(*alphabet*) = *a*). The mapping between *C*_*SAX,j*_ and 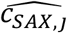 is obtained as follows: 
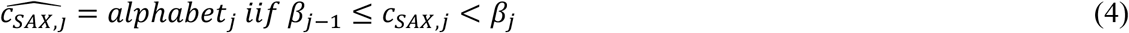

The set *β* = {*β*_*j*_, *j* ∈ ⟦1, *a* − 1⟧} is obtained by partitioning the set of values in equiprobable regions under a Gaussian distribution assumption.

There is no recommendation for choosing the parameters (*w*, *a*). To adopt a fully unsupervised approach, we chose the combination of parameters (*w*, *a*) that maximizes the average silhouette[28]. We applied agglomerative hierarchical clustering on the distances matrix computed. We chose the number of clusters using the Silhouette method[29].

### 2.5 Statistical Software, Reproducibility and Ethical Considerations

All scripts are available at *https://github.com/equipe22/ICDtimeliness*. All analyses were performed with the R statistical software v.3.4.2. Temporal clustering was performed using the TSclust package[30]. This study was conducted under the methodological guidelines MR-005 of the French national data privacy authority[25].

## 3 RESULTS

### 3.1 Description of ICD codes

We extracted 238,334,751 encounters between 2008, January 1^st^ and 2017, December 31^st^, corresponding to 56,621,773 distinct patients. The number of encounters per month slightly increased from 1,842,241 in January 2008 to 2,156,443 in December 2017. The number of patients per month slightly increased from 1,403,779 in January 2008 to 1,513,826 in December 2017. 1,006 ICD codes presented a variation of the relative amplitude of frequencies lower than 50%, 510 codes between 50% and 100% and 521 greater than 100%.

### 3.2 Temporal Clustering

Out of the 2,037 codes included in the study, we kept the 1,758 codes covering the entire period 2008-2017 for the clustering step. We extracted the moving means for the 1,758 ICD codes respecting the inclusion criteria. We identified the parameters of the SAX representation maximizing the silhouette: 20 frames (w), and alphabet size (a) of 6. The optimal number of clusters with the Silhouette and Elbow methods was 4. Figure 1 presents the dendrogram of the hierarchical clustering. Figure 2 presents the normalized moving mean time series for each cluster.

**Figure 1:**
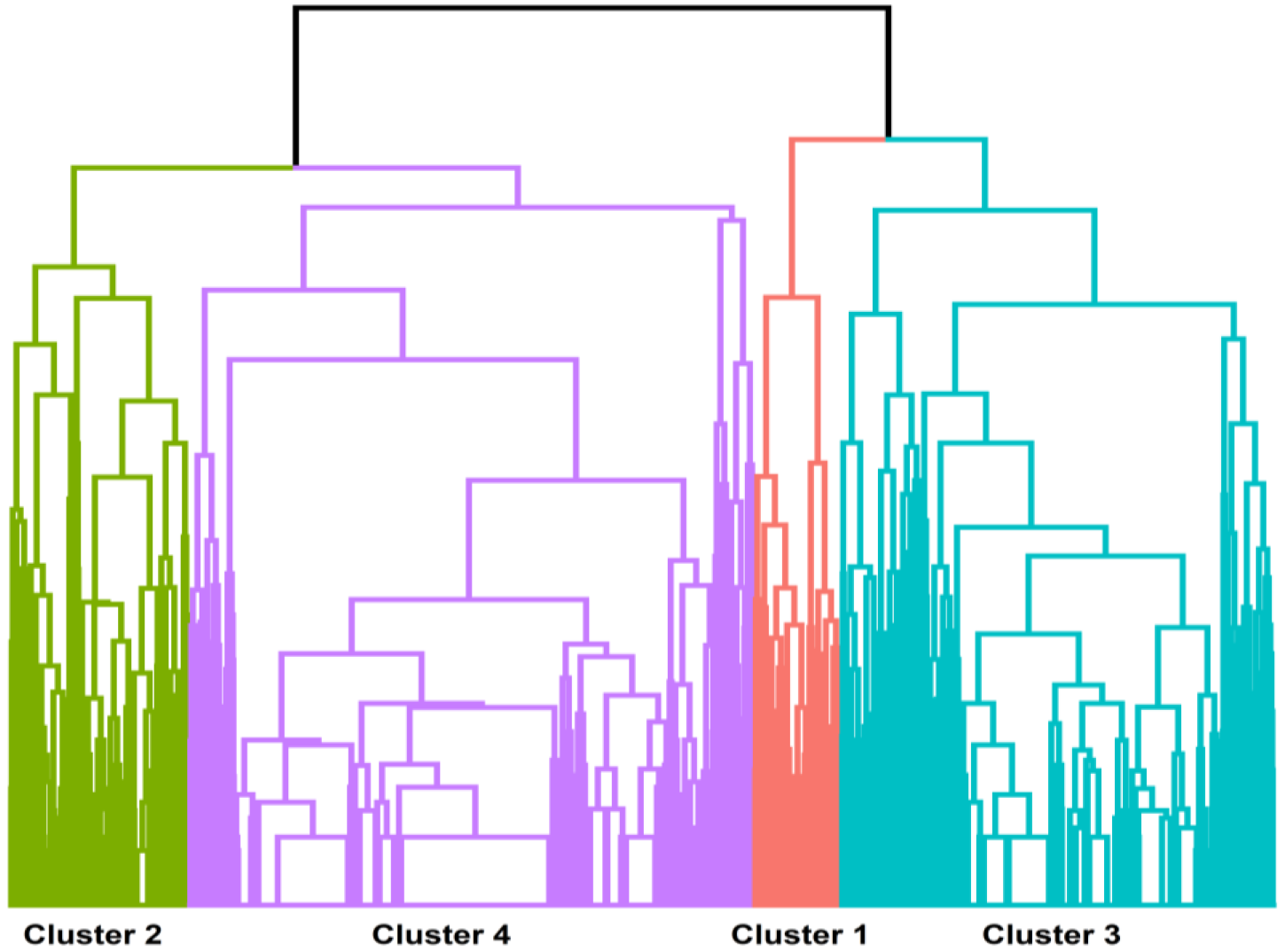
Dendrogram of the hierarchical clustering of the ICD normalized moving mean time series with the 4 selected clusters

**Figure 2:**
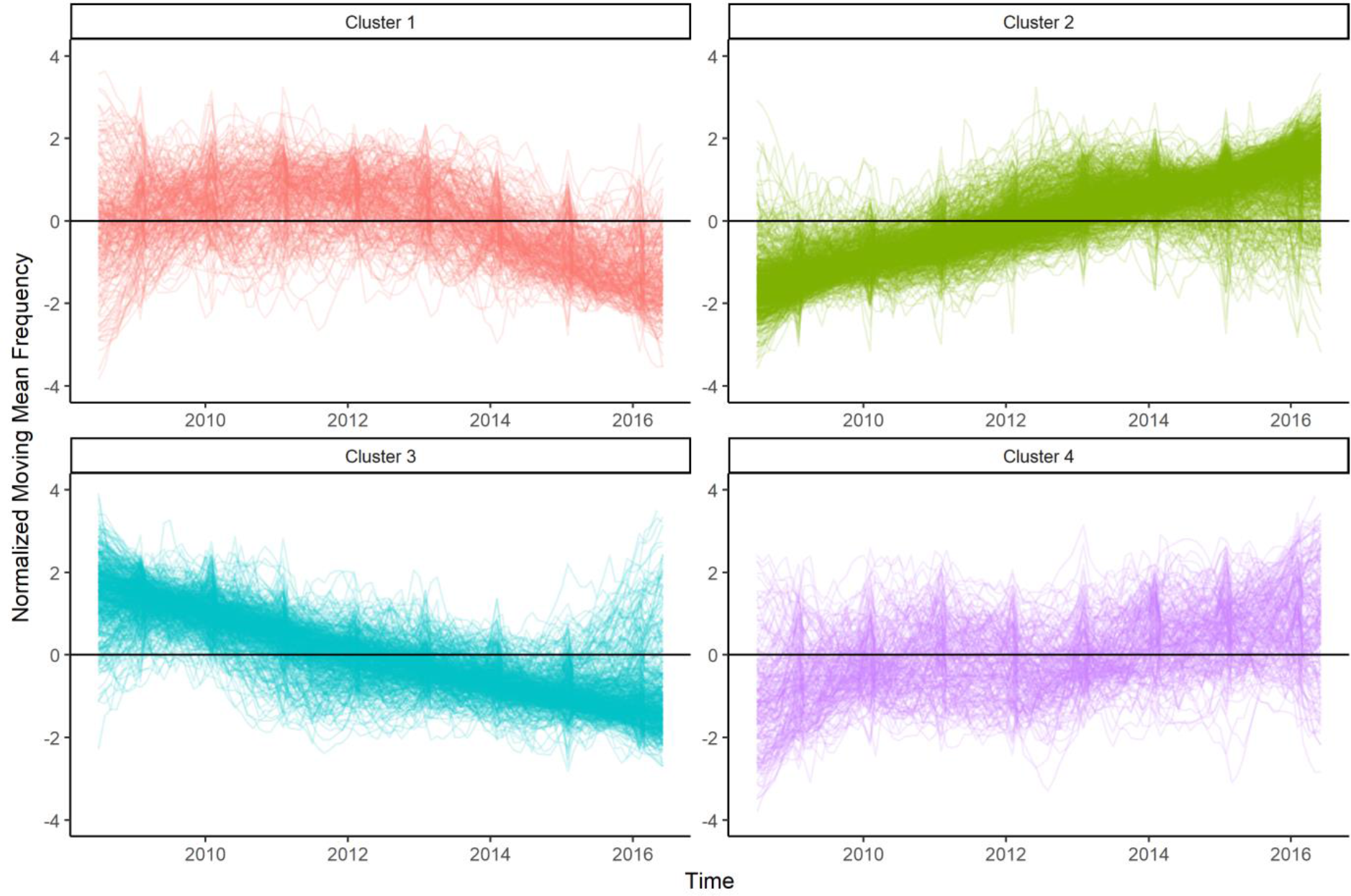
Representation of the ICD normalized moving mean time series by cluster

**Table 1:**
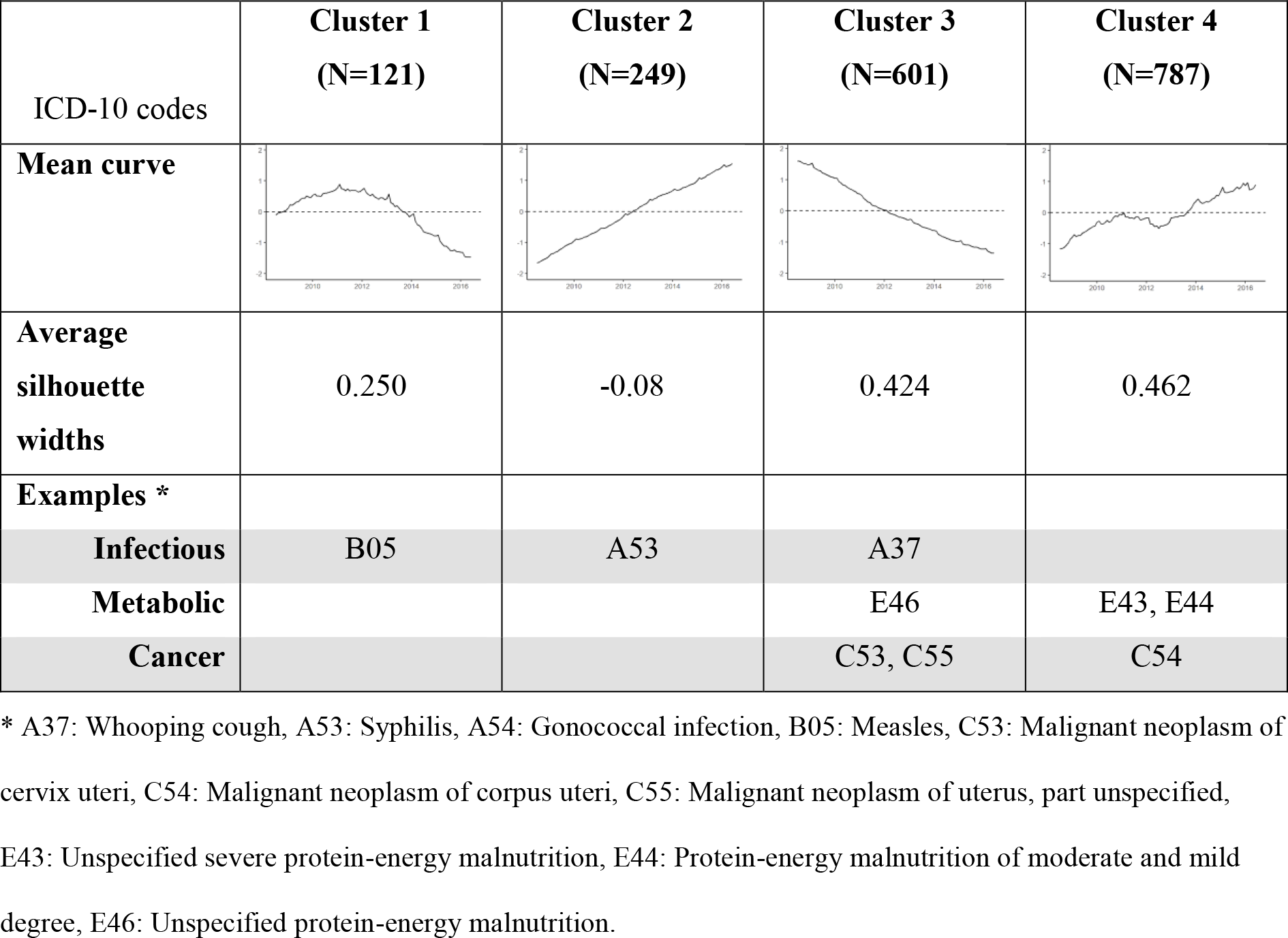
Characteristics of clusters and examples of codes.

| ICD-10 codes | Cluster 1<br>(N=121) | Cluster 2<br>(N=249) | Cluster 3<br>(N=601) | Cluster 4<br>(N=787) |
| --- | --- | --- | --- | --- |
| **Mean curve** |  |  |  |  |
| **Average silhouette widths** | 0.250 | -0.08 | 0.424 | 0.462 |
| **Examples \*** |  |  |  |  |
| **Infectious** | B05 | A53 | A37 |  |
| **Metabolic** |  |  | E46 | E43, E44 |
| **Cancer** |  |  | C53, C55 | C54 |

\* A37: Whooping cough, A53: Syphilis, A54: Gonococcal infection, B05: Measles, C53: Malignant neoplasm of cervix uteri, C54: Malignant neoplasm of corpus uteri, C55: Malignant neoplasm of uterus, part unspecified, E43: Unspecified severe protein-energy malnutrition, E44: Protein-energy malnutrition of moderate and mild degree, E46: Unspecified protein-energy malnutrition.

### 3.3 Examples of evolution profiles

We selected ICD codes in each cluster to illustrate the evolution profiles. Figure 3 presents the evolution profiles of ICD codes for infectious diseases. Figure 4 presents the evolution profiles of metabolic disease and cancer codes.

**Figure 3:**
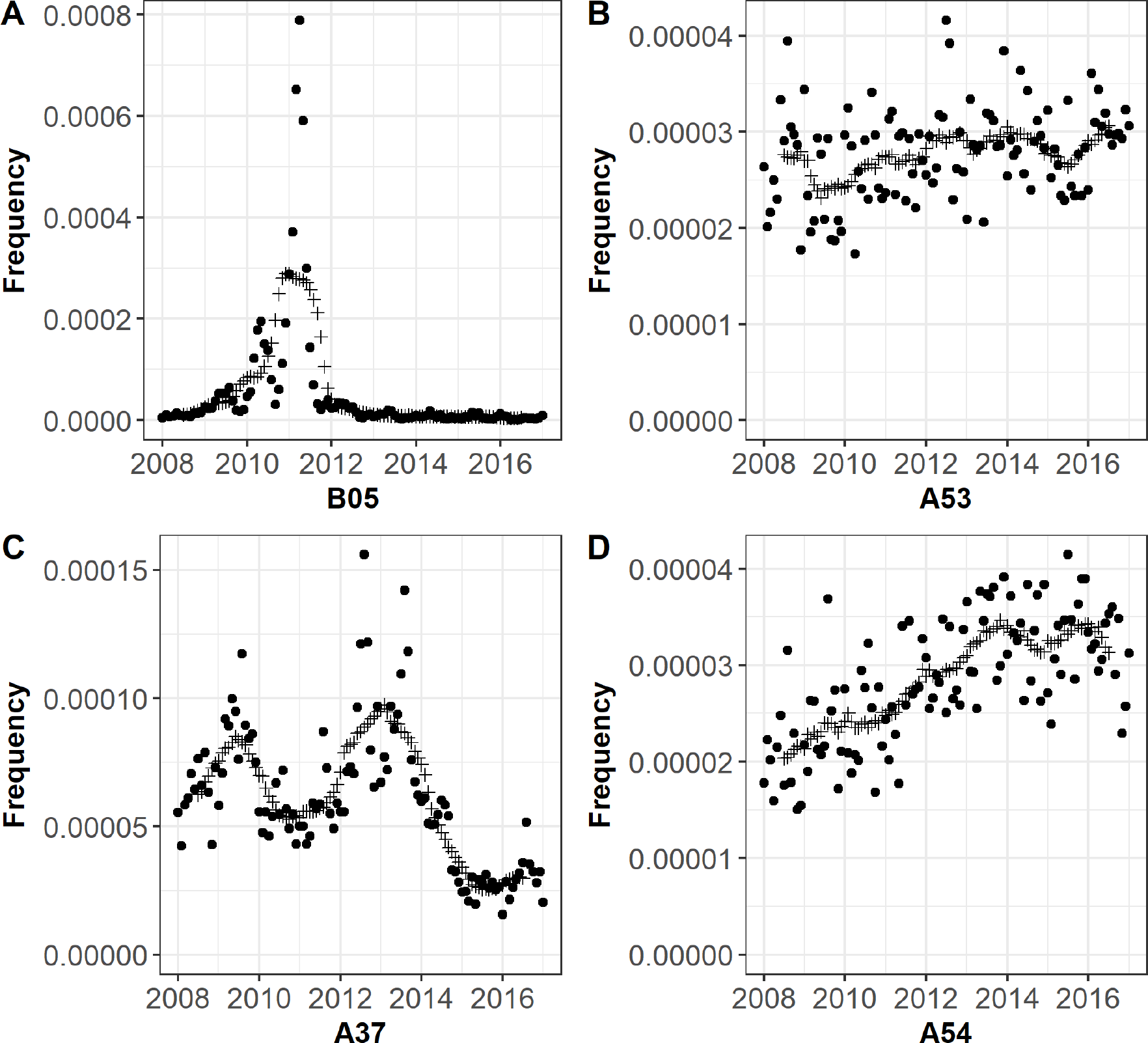
Evolution’s profiles of examples of infectious codes. A37: Whooping cough, A53: Syphilis, A54: Gonococcal infection, B05: Measles. Circles represent the actual values, plus a smoothed value over 12 months.

**Figure 4:**
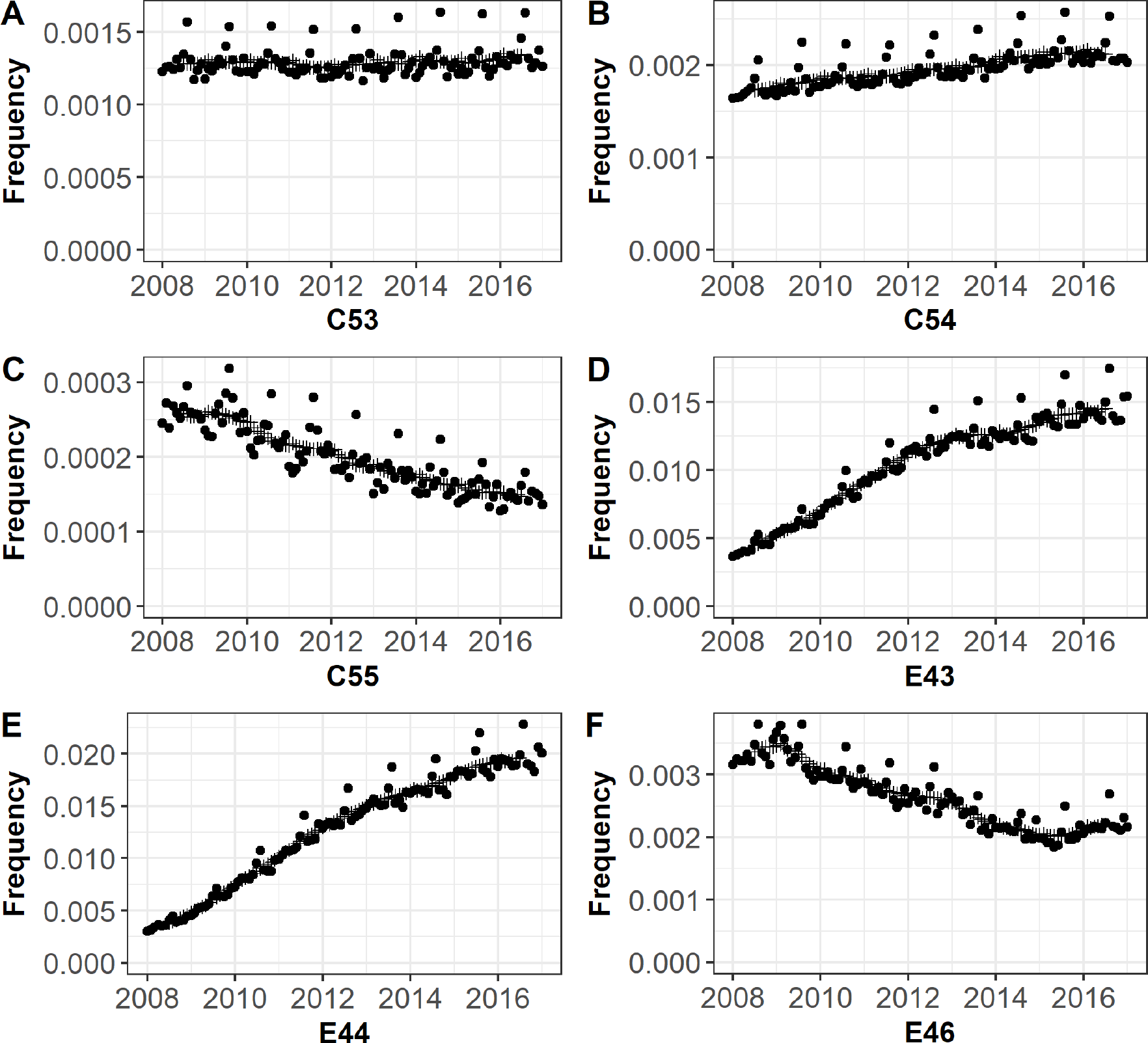
Evolution profiles of examples of metabolic and cancer codes. C53: Malignant neoplasm of cervix uteri, C54: Malignant neoplasm of corpus uteri, C55: Malignant neoplasm of uterus, part unspecified, E43: Unspecified severe protein-energy malnutrition, E44: Protein-energy malnutrition of moderate and mild degree, E46: Unspecified protein-energy malnutrition. Circles represent the actual values, plus a smoothed value over 12 months.

## 4 DISCUSSION

### 4.1 Technical Findings

The overall results showed a strong instability (i.e. large variation of frequency over time) of the use of ICD codes over time, with an important variation of the relative amplitude of the frequencies: 510 codes have relative variations comprised between 50% and 100% and 521 greater than 100%. We observed 784 codes with increasing profiles (cluster 4) and 601 codes with decreasing profiles (cluster 3). We analyzed the evolution of three categories of ICD codes (infectious, metabolic and cancer). We described the evolution of ICD codes of infectious diseases to valid our methodology. Infectious diseases are monitored through epidemiological surveillance networks and the outbreaks are measured. In appendices, we detail how examples presented in figure 3 are related to epidemiological phenomena. Epidemiological outbreaks are detected with our 6-months smoothing. However, all trends cannot be clearly explained by epidemiological events.

#### External and intrinsic factors

We distinguished between external factors due to changes in billing, organization, policy or regulation and intrinsic factors due to epidemiological phenomenon. External factors can have a major influence and bias the results of longitudinal studies. In appendice, we explored two cases of such phenomena: the increase of malnutrition ICD codes (E43, E44 and E46) explained by the evolution of practices of coding optimization (for financial reason, an example of billing effect); and the evolution of codes of malignant neoplasms of the uterus explained by an improvement of precision of coding (learning effect). The billing effect is a change of coding practice due to financial incitation of the national reimbursement policy. The learning effect is an improvement of the precision of coding due to a better knowledge of the coding system by the physicians.

### 4.2 Significance for Data Reuse

#### International relevance

This study focuses on a national database in France. Other national databases could reflect similar external and intrinsic evolution. Integrated health information systems are being developed for both public health and research purposes[31]. Collaborative studies across data sources in several countries require explicit documentation of the external factors that may influence the distributions of the variables stored in the distributed databases. Our result suggests that a common terminology does not guarantee interoperability. Interoperability is affected by many factors including the potential link between reimbursement policy and coding system[32] or the change in coding system policy[33].

Analytic interoperability has to move beyond the traditional mapping approaches, where the emphasis is exclusively on aligning database schemas and codes and implement solutions in which the analysis of data can be ported between data sets, where although the codes may have been mapped the characteristics of the populations may lead to incompatible results if the same algorithms or statistical analyses are naively run ‘as is’ across datasets. As more research projects and surveillance methods will rely on reusing health administrative databases, these efforts will become crucial.

#### Impact on phenotyping

Most of the studies looking for associations between genotypes and phenotypes reuse ICD codes from EHRs as phenotype information[7]. The timeliness issue may have an impact on phenotyping algorithms and analyses distributed through networks of clinical data warehouses. Phenotype algorithm use structured and unstructured data to better identify cohorts of subjects within the health data. ICD codes are often an important source of structured data. For example, in PheKB[34], 70% of the phenotype algorithms use ICD codes. Therefore, our results suggest that developing phenotyping algorithms from administrative database alone, without taking the context of evolution of the code, can lead to substantial measurement bias in population studies.

### 4.3 Related Work

This study takes place in the context of temporal plausibility. The general topic of data quality has been widely explored in the literature[35,36]. Kahn *et al.*[24] have defined the plausibility as “features that describe the believability or truthfulness of data values.” Kahn proposed a distinction between atemporal plausibility and temporal plausibility. The atemporal plausibility “seeks to determine if observed data values, distributions, or density agree with local or “common” knowledge (Verification) or from comparisons with external sources that are deemed to be trusted or relative gold standards (Validation)”. Whereas the temporal plausibility “seeks to determine if time-varying variables change values as expected based on known temporal properties or across one or more external comparators or gold standards.” Previous studies evaluated the quality of administrative databases in terms of validation studies for atemporal plausibility, e.g.,[37,38]. Quan *et al.*[37] studied the agreement between charts and ICD-9 codes on occurrence of procedures in patients admitted to either general medicine or general surgery services. They noticed a good specificity (greater than 99%) and an uncertain sensitivity (from 0 to 87.5%). They also analyzed positive predictive values, negative predictive values and kappa. This approach is classical in data quality analyses of administrative database. Hinds *et al.*[38] proposed a review of data quality studies of administrative health databases in Canada. Most of the data quality measures concern sensitivity (64.2%), specificity (55%), negative predictive value (43%), positive predictive value (58.3%) and kappa (30.5%). The coding errors were explained by various factors, such as clinicians' knowledge of terminologies, clinicians' experience in coding, diligence for compiling information, transcriber's ability to read notes and EHRs, and even intentional errors like underspecification and upcoding[39].

### 4.4 Remaining Challenges

#### Clusters and exploration of internal and external causes

In our study, we have adopted a fully unsupervised approach for temporal clustering to explore the existence of evolution profiles without prior knowledge on the evolution. Other approaches have been developed, for example Sacchi *et al.*[40] proposed a two-step approach combining a qualitative representation of time series and a hierarchical clustering. A parametric approach based on piecewise regressions could provide another description for hypothesis-driven exploration. The main challenge resides in the identification of the origin of the evolution (intrinsic epidemiological or external factors). The detailed analysis of profiles show that the same cluster can contain profiles influenced by either cause and in some case, both causes simultaneously. In most cases, additional knowledge and sources are probably required to determine automatically the origin of the profile.

#### Contextualization

Saez *et al.*[22] have defined the contextualization as “the degree to which data is correctly/optimally annotated with the context in with it was acquired.” We measured the stability of the frequencies and proposed a contextualization for a limited number of ICD-10 codes. The retrospective annotation of the timeliness of all the ICD codes is an important task and a perspective for a future work. We proposed some fundamental dimensions of the timeliness assessment. Other DQ metrics can be applied as the Information-Geometric Temporal or the Probability distribution function statistical process control[41]. Furthermore, we limited our analysis to the observation of the timeliness dimension. The DQ assessment of administrative database is broader than the timeliness dimension, for example, the national hospitalization database can be seen as multisources data and other DQ metrics like the source probabilistic outlyingness metric or the global probabilistic deviation metric could be used[42].

### 4.5 Recommendations

In order to address the different aspects of analytic harmonization, the health research community must develop a set of metadata. These metadata should represent this information in such a way as to facilitate comparison between data sources enabling the translation of research questions between these sources. While research efforts have developed shared frameworks for provenance[43,44], our study demonstrates that we still need a framework for the capturing and representation of information about data quality. Data quality is a shared concern in research, and in public health[45]. Therefore, data quality profiling as metadata should be integrated in big data infrastructure as proposed by Merino *et al.*[46] in their “Data-Quality-in-Use model.” Metadata definitions and ontologies for data sharing will enable the fingerprinting of data repositories and make the researchers aware of the quality of the available data. An IT framework for capturing detailed formal metadata about data sources, based on shared catalogs and automated data annotation would facilitate systematic annotation and harmonization.

## 5. CONCLUSION

We proposed an evaluation of the timeliness of ICD-10 codes in the French hospitalization summary discharges. External and intrinsic factors influencing the data quality of administrative data are heterogeneous and time-varying; human annotation remains necessary. A key challenge is to annotate these data repositories by producing machine-readable metadata to foster the improvement of the machine learning performances.

## APPENDICE: Looking for Extrinsic and Intrinsic Factors Explaining the Observed Evolution of the Examples of Figures 3 and 4

***Figure 3A (B05, Measles).*** The prevalence of measles is close to zero and stable over time, except around 2011 where an explosive peak is detected. More precisely, we observed three peaks of increasing intensity between 2009 and 2012. As measles is a notifiable disease in France, cases are exhaustively reported in national registers. Based on these national surveillance data, Antona *et al.*[1,2] reported that France experienced a massive measles outbreak in 2010-2011, accounting for more than half of the 30,000 cases in Europe during this period, with almost 5,000 persons hospitalized. The epidemic curve of their article showed that the number of cases started increasing in mid-2008, evolving in 3 epidemic waves: 2009, 2010 and the highest epidemic waves in 2011. Our results are consistent with this data.

***Figure 3B (A53, Syphilis).*** The prevalence of syphilis slightly increased since 2010. In France, the *RésIST* network contributes to monitoring sexually transmitted infections such as Syphilis or Gonorrhea. The collection of demographic, clinical, biological and behavioral data is based on different diagnostics, information and screening sites, and medical consultations.

Surveillance data from RésIST [3] confirmed our observed trend, which translates the increase in risky sexual behaviors, in particular in men who have sex with men. Despite the heterogeneity of surveillance systems, European surveillance data lead to the same findings [4].

***Figure 3C (A37, Whooping cough).*** On Figure 3C, we observe two outbreaks in 2009 and 2012. After 2012, the prevalence of whooping cough decreased until 2016. In France, *RENACOQ*, a sentinel hospital-based voluntary surveillance network has been established in 1996, and covers about 30% of hospitalized pertussis pediatric cases. Surveillance data from RENACOQ [5,6] confirmed our observed outbreaks: there have been increases in pertussis prevalence in the past few years in France, and in particular, two epidemic peaks occurred in 2009 and 2012.

***Figure 3D (A54, Gonococcal infection).*** Gonococcal infections highly increased between 2008 to 2017. Surveillance data from RésIST, and Renago, a laboratory network that collects demographic and biological data for gonorrhea, show similar results. The diffusion of multi-resistant strains in a context of more and more frequent transmission of gonococci explains these trends[7,8].

***Figures 4A, 4B and 4C (C53, Malignant neoplasm of cervix uteri; C54, Malignant neoplasm of corpus uteri; C55, Malignant neoplasm of uterus, part unspecified).*** The C53 code (Malignant neoplasm of cervix uteri) showed a stationary trend whereas French and European literature data reported a decreasing incidence over the past years, mainly due to the widespread implementation of screening programs (cervical smear tests) since the 1960s and HPV vaccination since 2006 in France[9–11]. Conversely, the C54 code (Malignant neoplasm of corpus uteri) increase slightly during the study period, in accordance with epidemiological data [9,10,12]. The unspecified code C55 (Malignant neoplasm of uterus, part unspecified) shows a 2-fold decrease between 2008 and 2017. By analyzing the curves together, we can suppose that the stationary trend of C53 may be explained by an improvement of coding precision, leading to abandon the C55 code. This billing effect masked the real cervix cancer evolution over time. To analyze time trends of uterine cancer incidence with data extracted from the ICD-10 classification, we had to deal with varying coding precision over time. Loos *et al.* focused on imprecisely coded uterine cancer deaths based on ICD-10 codes and developed a reallocation procedure for the “unspecified’ category. This method may be added as a supplementary step in incidence data treatment based on ICD codes to avoid misleading interpretation[13].

***Figures 4D, 4E and 4F (E43, Unspecified severe protein-energy malnutrition; E44, Protein-energy malnutrition of moderate and mild degree; E46, Unspecified protein-energy malnutrition).*** In figure 4D, 4E, 4F we explored the E43, E44, E46 malnutrition codes. These codes are often used in France to demonstrate the severity of a hospital stay and their coding is therefore used to optimize reimbursement by hospitals. The E43 code (Unspecified severe protein-energy malnutrition) and the E44 code (Protein-energy malnutrition of moderate and mild degree) show a global increase. Considering these results, malnutrition during hospital stays would have increased by a factor of 6 between 2008 and 2017. There are no available arguments that can justify these trends. Moreover, undercoding of malnutrition has been highlighted in France as early as 2003[14]. In recent years, practices of coding optimization have been introduced in many institutions[15,16]. Therefore, coding optimization may explain the decreasing trend of E46 (Unspecified protein-energy malnutrition) for the benefit to E43 and E44 codes that entailed a gain of precision coding. The figure 4D (E43) presents a reduced slope starting in 2012, probably because financial optimization was reached globally, and no further improvement could be achieved.

## ABBREVIATIONS

DQ: Data Quality
EHR: Electronic Health Records
ICD: International Statistical Classification of Diseases and Related Health Problems
OHDSI: Observational Health Data Sciences and Informatics
SAX: Symbolic Aggregate Approximation representation
SNDS: *Système National des Données de Santé* (National Health Data System)

